# Expression of mammalian Onzin and Fungal Cadmium Resistance 1 in S. *cerevisiae* suggests ancestral functions of PLAC8 proteins in regulating mitochondrial metabolism and DNA damage repair

**DOI:** 10.1101/403675

**Authors:** Stefania Daghino, Luigi Di Vietro, Luca Petiti, Elena Martino, Cristina Dallabona, Tiziana Lodi, Silvia Perotto

## Abstract

Protein domains are structurally and functionally distinct units responsible for particular protein functions or interactions. Although protein domains contribute to the overall protein function(s) and can be used for protein classification, about 20% of protein domains are currently annotated as “domains of an unknown function” (DUFs). DUF 614, a cysteine-rich domain better known as PLAC8 (Placenta-Specific Gene 8), occurs in proteins found in the majority of Eukaryotes. PLAC8-containing proteins play important yet diverse roles in different organisms, such as control of cell proliferation in animals and plants or heavy metal resistance in plants and fungi. For example, Onzin from *Mus musculus* is a key regulator of cell proliferation, whereas FCR1 from the ascomycete *Oidiodendron maius* confers cadmium resistance. Onzin and FCR1 are small, single-domain PLAC8 proteins and we hypothesized that, despite their apparently different role, a common molecular function of these proteins may be linked to the PLAC8 domain. To address this hypothesis, we compared these two PLAC8-containing proteins by heterologous expression in the PLAC8-free yeast *Saccharomyces cerevisiae*. When expressed in yeast, both Onzin and FCR1 improved cadmium resistance, reduced cadmium-induced DNA mutagenesis, localized in the nucleus and induced similar transcriptional changes. Our results support the hypothesis of a common ancestral function of the PLAC8 domain that may link some mitochondrial biosynthetic pathways (i.e. leucine biosynthesis and Fe-S cluster biogenesis) with the control of DNA damage, thus opening new perspectives to understand the role of this protein domain in the cellular biology of Eukaryotes.

**Author Summary:** Protein domains are the functional units of proteins and typically have distinct structure and function. However, many widely distributed protein domains are currently annotated as “domains of unknown function” (DUFs). We have focused on DUF 614, a protein domain found in many Eukaryotes and better known as PLAC8 (Placenta-Specific Gene 8). The functional role of DUF 614 is unclear because PLAC8 proteins seem to play important yet different roles in taxonomically distant organisms such as animals, plants and fungi. We used *S. cerevisiae* to test whether these apparently different functions, namely in cell proliferation and metal tolerance, respectively reported for the murine Onzin and the fungal FCR1, are mediated by the same molecular mechanisms. Our data demonstrate that the two PLAC8 proteins induced the same growth phenotype and transcriptional changes in *S. cerevisiae*. In particular, they both induced the biosynthesis of the amino acid leucine and of the iron-sulfur cluster, one of the most ancient protein cofactors. These similarities support the hypothesis of an ancestral function of the DUF 164 domain, whereas the transcriptomic data open new perspectives to understand the role of PLAC8-proteins in Eukaryotes.

## Introduction

The PLAC8 domain was described for the first time in the protein Onzin, the product of the human Placenta-Specific Gene 8 [1]. The same domain was later identified in many eukaryotes, but its biological role remains elusive because PLAC8 proteins seem to play diverse roles in different organisms and cell types. The mammalian Onzin has been reported as a repressed target of the c-Myc oncoprotein [2], with pro-proliferative anti-apoptotic effects in many cell types and a role in leukemia [3], hepatic, pancreatic [4, 5] and colon cancer progression [6, 7], but also in adipocyte growth [8]. The same protein has pro-apoptotic activity in other cell types [9], indicating that the final effect of Onzin is highly dependent on cell type. In plants, PLAC8 genes are also involved in cell proliferation because mutants are altered in organs size. The tomato Fruit Weight 2.2 (FW2.2) negatively influences fruit size and is downregulated in domesticated species [10]. Similar function has been reported for FW2.2-like genes in other plant species, both dicots [11-14] and monocots [15,16].

Another cellular role of PLAC8 proteins in plants and fungi is to increase resistance to heavy metals. First described in *Arabidopsis thaliana*, the Plant Cadmium Resistance (PCR) protein family includes proteins that confer resistance to cadmium or zinc. AtPCR1 was suggested to be - or to be part of-a cadmium transporter because of its localization on the plasma membrane in both *A. thaliana* and *S. cerevisiae* [17], while AtPCR2 was associated to zinc transport [18]. In fungi, the only PLAC8 genes characterized to date are two Fungal Cadmium Resistance (FCR) genes identified in a metal tolerant isolate of the mycorrhizal ascomycete *Oidiodendron maius* [19, 20], which increased cadmium resistance when expressed in *S. cerevisiae*. In particular, OmFCR1 (hereafter FCR1) localized in the yeast nucleus and physically interacted with Mlh3p, a key player in meiotic crossing-over and a subunit of the DNA mismatch repair (MMR) complex [19]. Despite the numerous studies on PLAC8 proteins in different organisms, it is still unclear whether the PLAC8 domain has a molecular function common to all biological systems. To address this question, we have used *S. cerevisiae* as a model organism to investigate two small, single-domain PLAC8 proteins from taxonomically distant organisms, the mammalian Onzin from *Mus musculus* and the fungal FCR1 from *O. maius*. *S. cerevisiae* is a good model system because it has been successfully used to test plant and fungal PLAC8 gene functions by heterologous expression [17-20]. In addition, although *S. cerevisiae* lacks genes coding for PLAC8 proteins this protein domain can be found in the genome of some basal fungi and other members of Saccharomycotina (see Mycocosm, https://genome.jgi.doe.gov/programs/fungi/index.jsf), thus suggesting that the metabolic framework interacting with this protein domain is found early in fungal evolution. *S. cerevisiae* genome lacks PLAC8 genes, but it

When expressed in cadmium-exposed *S. cerevisiae*, Onzin and FCR1 displayed very similar phenotypes, as they both conferred cadmium resistance and increased cell proliferation and vitality. Both proteins localized in the yeast nucleus, reduced the mutation frequency in homonucleotide runs and determined similar transcriptomic changes in cells exposed to cadmium. In particular, both PLAC8 proteins up-regulated ancient and conserved metabolic pathways that link mitochondrial functions related to leucine biosynthesis, Fe-S cluster biogenesis and maintenance of nuclear DNA integrity. Although the exact role of the PLAC8 domain remains unclear, our findings provide support to the hypothesis of a common ancestral function for this protein domain.

## Results and discussion

Onzin is a small protein highly conserved in all vertebrates and in particular in mammals, where mouse and human orthologous proteins are 83% identical (Fig 1). By contrast, Onzin shared only 29 amino acids with the fungal PLAC8 protein FCR1, with an overall 25% sequence identity mainly confined to the signature cysteine-rich motif of the PLAC8 domain (Fig 1).

**Fig 1.**
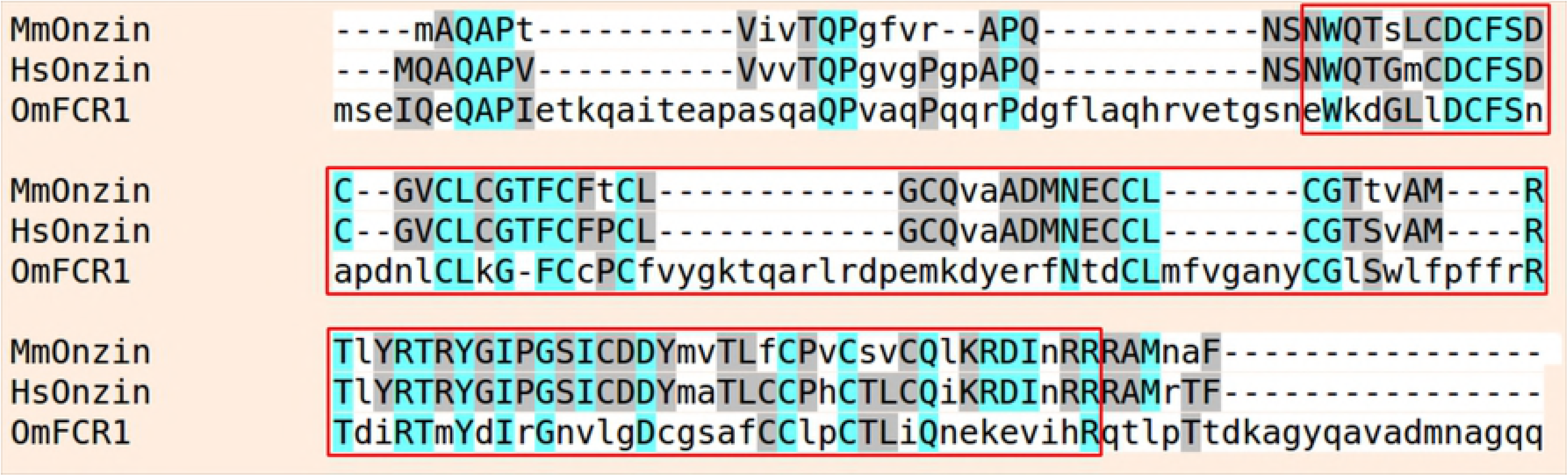
Amino acid sequence alignment of PLAC8 domain-containing proteins. The complete sequences of MmOnzin (from *Mus musculus*), HsOnzin (from *Homo sapiens*) and OmFCR1 (from *Oidiodendron maius*) have been aligned. Similar residues are colored as the most conserved one according to BLOSUM62 average scores: Max: 3.0 (light blue), Low: 0.5 (grey). Lower case non-colored letters indicate amino acid residues with no similarities. The protein alignment was performed using the Phylogeny.fr platform. The red box shows the PLAC8 domain.

A first set of experiments was aimed to verify whether Onzin expression in cadmium-exposed *S. cerevisiae* induced a phenotype similar to the one previously described for FCR1 [19]. Spot dilution assays at two CdSO_4_ concentrations (Fig 2A) showed that both Onzin and FCR1 confer cadmium tolerance to *S. cerevisiae*, when compared with the control strain transformed with the empty vector. At 25 μM CdSO_4_, Onzin-expressing yeast grew even better than the FCR1-expressing strain (Fig 2A). A truncated Onzin lacking the conserved N-terminal region of the PLAC8 domain (Onzin^Δ28-38^) led to a partial loss-of-function phenotype (Fig 2A), suggesting that this protein region is important for Onzin function. Evaluation of the cadmium half inhibitory concentration (IC50) on the same yeast strains grown in liquid culture yielded fully consistent results (S1 Fig).

**Fig 2.**
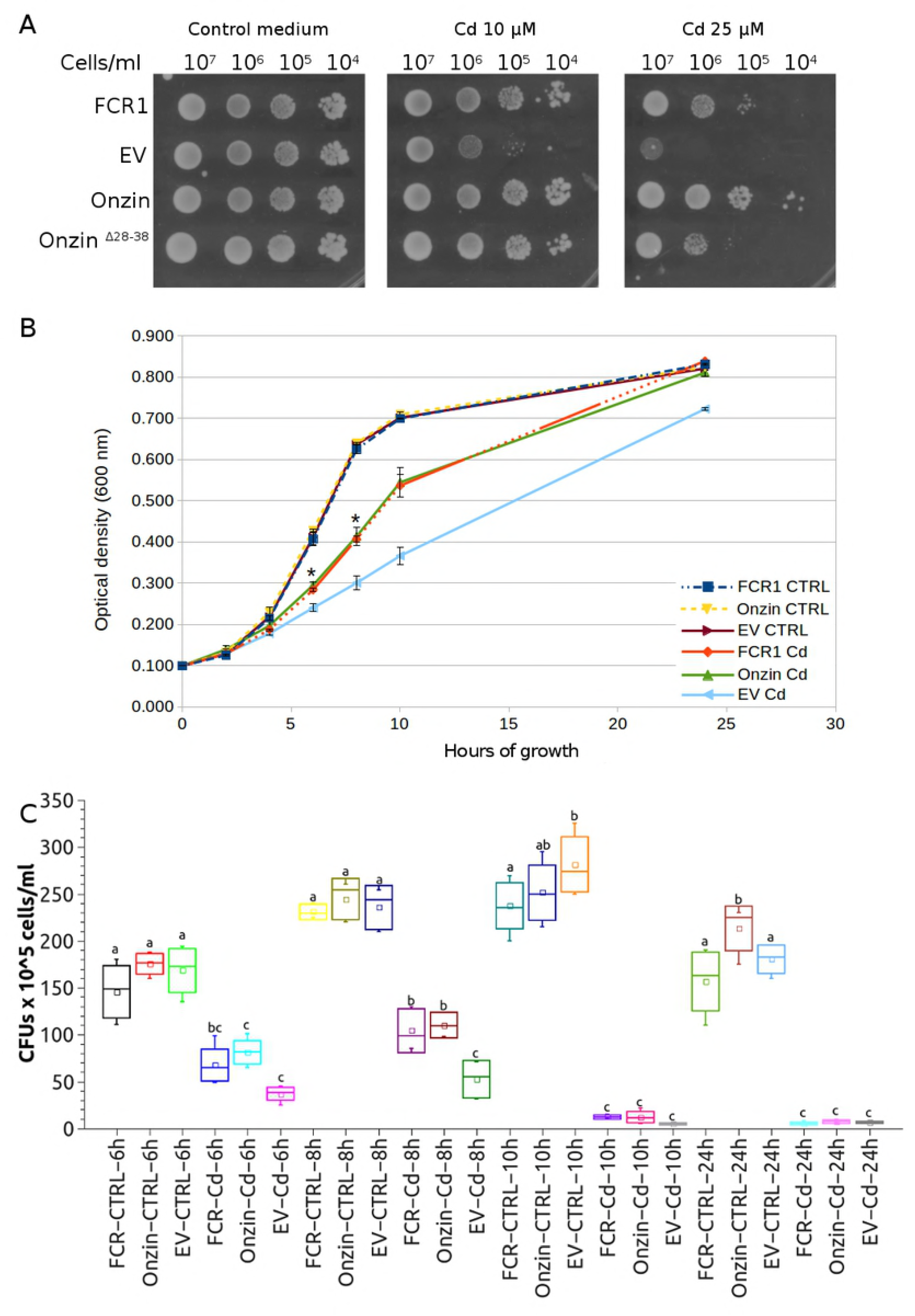
Growth and viability of yeast cells expressing FCR1 and Onzin on cadmium-containing media. (A) Spot dilution assay of yeast (EAY1269 strain) expressing FCR1, wild-type Onzin, the truncated Onzin^Δ28-38^ or the empty vector pFL61 (EV). Strains were plated in ten-fold serial dilutions onto YNB-D medium amended with CdSO_4_ (10 or 25 µM) or not (control medium). (B) Growth curves of yeast cell cultures in control medium (CTRL) or in a medium containing 25 µM CdSO_4_ (Cd). The optical density (OD_600_) of cultures expressing FCR1, Onzin and pFL61 (EV) was measured after 2-4-6-8-10-24 hours of incubation at 30°C and 150 rpm. The asterisks indicate time points with a significant difference (n=3, Shapiro Wilk as normality test, ANOVA with Tukey P<0.01) between cells expressing FCR1, or Onzin, and cells transformed with the empty vector (EV). (C) Cell viability of yeasts expressing FCR1, Onzin or the empty vector pFL61 (EV) after growth for 6- 8-10-24 hours in control medium (CTRL) or in a medium containing 25 µM CdSO_4_ (Cd). Colony Forming Units (CFUs) were counted for each yeast culture. Samples showing statistically different CFU numbers (P<0.05 by ANOVA with Tukey as post-hoc test, n=6 for 6-10-24h time points, n=3 for the 8h time point, Shapiro Wilk as normality test) are indicated by different letters. The square symbol indicates the mean value. The wiskers indicate the minimum and the maximum values. The top and the bottom of the rectangle indicate ± standard deviation, while the central line of the rectangle indicates the 50%.

Yeast growth and viability was also monitored for 24 h in liquid culture on control medium and on medium containing 25μM CdSO_4_. On control medium, all yeasts strains showed identical growth curves, as measured by OD_600_, whereas on cadmium-amended medium Onzin and FCR1 expressing yeasts grew more than the control strain, starting from 6 h incubation (Fig 2B). Cell survival in the same growth experiment was measured by colony forming units (CFUs) count and was similar for all yeast strains grown on control medium. By contrast, Onzin and FCR1 expression led to CFU numbers higher than the empty vector after 6 and 8 h of cadmium exposure (Fig 2C). At later time points, cell survival on cadmium-amended medium was low for all yeast strains (Fig 2C).

Subcellular Onzin localization (Fig 3) in yeast cells by C-terminal tagging with the Enhanced Green Fluorescent Protein (EGFP) showed that Onzin-EGFP co-localized, like FCR1-EGFP, with the Red Fluorescent Protein (RFP) fused to the nuclear localization signal (NLS).

**Fig 3.**
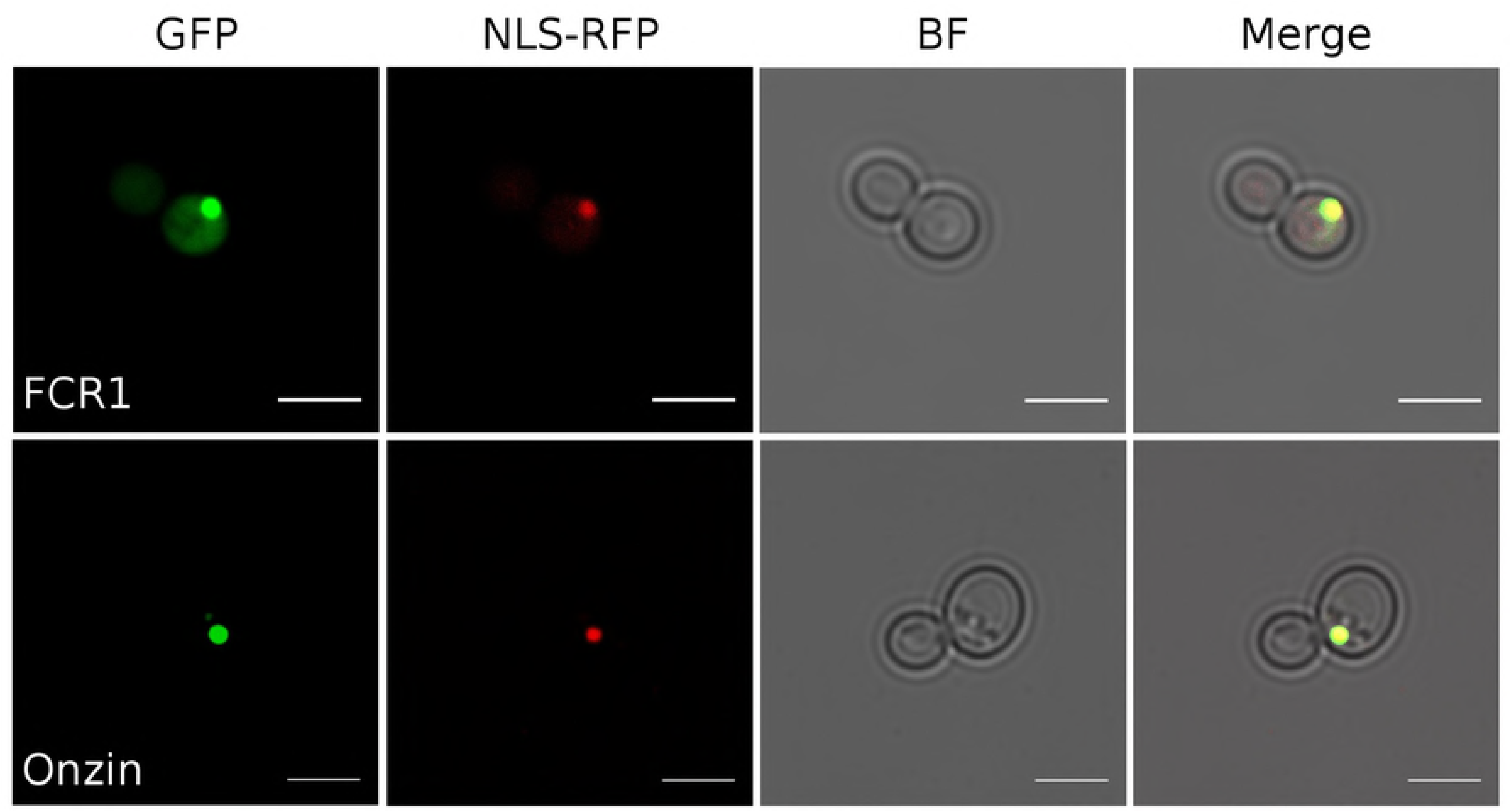
Subcellular localization of FCR1 and Onzin proteins in yeast cells. The FCR1-GFP and the Onzin-GFP fusion proteins were localized to the yeast nucleus, as indicated by co-localization (Merge) with a fusion protein carrying a nuclear localization signal (NLS)-RFP. BF: bright field image of the yeast cells. Scale bar is 5μm.

Thus, this first set of experiments showed, for Onzin-expressing yeast, the same phenotype already described for FCR1 [19]. FCR1 and Onzin expression increased yeast growth exclusively on cadmium-containing medium, indicating a protein function only measurable during cadmium stress. The nuclear localization of EGFP-tagged FCR1 and Onzin would exclude a direct role in metal detoxification mechanisms, such as membrane transport or scavenging. We therefore investigated possible functions of the two PLAC8 proteins correlated with their nuclear localization.

### Both PLAC8 proteins physically interact with Mlh3p and reduce cadmium induced DNA mutations

Cadmium does not damage DNA directly but is mutagenic because it interferes with the cellular response to DNA damage [21] and inhibits all major DNA repair pathways, including the mismatch repair (MMR) complex [21-23]. FCR1 was found to physically interact with Mlh3p, a component of the MMR complex [19] and a yeast two hybrid assay confirmed its interaction with the C-terminal region of both the yeast and the *O. maius* Mlh3p (S2 Fig). The same assay revealed a similar, albeit weaker, interaction of Onzin with the C-terminal region of the yeast and the mouse Mlh3 proteins (S2 Fig).

Previous experiments using the forward mutation assay at the canavanine-resistance (*CAN1*) locus did not reveal an influence of FCR1 on the DNA mutation rate under cadmium stress [19]. However, the *CAN1* assay could reveal only small differences between wild-type and a *mlh3*-defective yeast [24]. Here, we investigated the influence of FCR1 and Onzin expression on cadmium-induced mutagenicity with the more sensitive yeast *lys2*::*insE-A_14_* reversion assay, based on the restoration of the open reading frame in a mononucleotide run of 10 adenines within the *lys2::insE-A_14_* allele [25]. In cells defective of the MMR system, this assay could reveal 10-to 1,000-folds increase in mutations [26]. On control medium, yeast strains transformed with the empty vector or with the two PLAC8 genes showed no differences in DNA mutation rate (Table 1). As expected, cadmium exposure increased the DNA mutation rate, but expression of both FCR1 and Onzin reduced cadmium-induced DNA mutagenesis about three-folds when compared with the empty vector (Table 1). As FCR1 and Onzin physically interact with Mlh3p, a *mlh3* defective mutant was used to investigate whether reduction in the mutation rate required this protein. The *mlh3* strain transformed with the empty vector showed a higher background mutation rate both in control medium and in cadmium-amended medium, as already shown [23], but expression of FCR1 and Onzin led to a five-fold reduction in the DNA mutation rate when this mutant was exposed to cadmium (Table 1). Thus, yeast cells exposed to cadmium display a reduced DNA mutation rate when they express either FCR1 or Onzin, although the relationship between these two PLAC8 proteins and the MMR complex remains unclear.

**Table 1.** Influence of Onzin and FCR1 on DNA mutation rate measured by the Lys2::insE-A14 reversion assay.

| Strain | Vector | Treatment | Reversion Rate x 10<sup>-5</sup><br>(confidence intervals) | Fold increase in<br>mutation rate<sup>a</sup> | n<sup>b</sup> |
| --- | --- | --- | --- | --- | --- |
| WT | pFL61-EV | Control medium | 0,14 (0,30÷0,09) | 1 | 30 |
|  | pFL61-FCR1 |  | 0,19 (0,21÷0,13) | 0,74 | 20 |
|  | pFL61-Onzin |  | 0,18 (0,20÷0,16) | 0,74 | 20 |
| WT | pFL61-EV | CdSO<sub>4</sub> 1μM | 5,49 (6,10÷5,00) | 1 | 30 |
|  | pFL61-FCR1 |  | 1,75 (1,88÷1,54) | 3,14 | 20 |
|  | pFL61-Onzin |  | 1,92 (1,99÷1,70) | 2,85 | 20 |
| mlh3 | pFL61-EV | Control medium | 1,55 (1,88÷1,03) | 1 | 25 |
|  | pFL61-FCR1 |  | 1,63 (1,80÷1,20) | 0,95 | 25 |
|  | pFL61-Onzin |  | 1,57 (1,68÷1,04) | 0,99 | 25 |
| mlh3 | pFL61-EV | CdSO<sub>4</sub> 1μM | 14,61 (15,11÷14,01) | 1 | 25 |
|  | pFL61-FCR1 |  | 2,8 (3,01÷2,03) | 5,21 | 25 |
|  | pFL61-Onzin |  | 2,55 (2,97÷2,12) | 5,72 | 25 |

Reversion to Lys^+^ phenotype was assessed in the wild-type (WT) and *mlh3* yeast mutant strains expressing FCR1, Onzin or the empty vector (EV) on both control medium and cadmium-containing medium. ‘n’ column indicates the number of independent cultures tested from at least two independently constructed strains. Median mutation rates are presented as x10^-5^ with 95% confidence intervals. Relative mutation rates are compared with the empty vector of the same genetic background strain. ^a^each median reversion rate was normalized to the empty vector median rate to calculate the fold increase; ^b^n=number of independent replicates.

### Onzin and FCR1 induce similar transcriptomic changes in cadmium-exposed yeast

To identify the yeast pathways transcriptionally regulated by the two PLAC8 proteins, an RNAseq experiment was performed after 8 h of exposure to 25 μM CdSO_4_. The diagram in S3 Fig reports the number of transcripts significantly regulated in yeast expressing either FCR1 or Onzin, as compared with the empty vector (log_2_ fold-change threshold >1 or <-1, adjusted p-value <0.05). Genes up-regulated by both proteins, indicated as *PLAC8 up-regulated* genes, represented 68% and 70% of the total number of genes up-regulated by FCR1 and Onzin, respectively. Genes down-regulated by both proteins, indicated as *PLAC8 down-regulated* genes, represented 59% and 72% of the total number of genes down-regulated by FCR1 and Onzin, respectively. A complete list of FCR1 and Onzin regulated genes is available in the S1 Table.

“Mitochondrion” and “mitochondrial parts” were the only Gene Ontology (GO) enriched cellular compartments identified among *PLAC8 up-regulated* genes, together with the biological processes involved in the biosynthesis of the branched-chain amino acids, and the metal ion binding molecular functions, in particular iron (S2 Table). Consistently with these GO data, *PLAC8-upregulated* genes were involved in several mitochondrial pathways. “Plasma membrane” was the cellular compartment enriched in *PLAC8 down-regulated* genes, together with ions, amino acids and sugars transport molecular functions, and iron homeostasis (S2 Table).

### Onzin and FCR1 do not activate antioxidant responses to cadmium

Activation of antioxidative enzymes and metabolites is a common cell defense response that could reduce cadmium toxicity [27]. Although cadmium is unable to generate free radicals directly, cadmium exposure induces in fact the production of reactive oxygen species (ROS). Activation of antioxidative enzymes and metabolites is therefore a common cell defense response that could reduce cadmium toxicity [27]. Previous experiments [19] suggested that FCR1 does not confer cadmium tolerance by increasing the antioxidative cell potential, and our transcriptomic data confirm this observation, as no genes coding for enzymes or compounds involved in ROS scavenging (e.g. superoxide dismutases or enzymes involved in glutathione metabolism) were identified among the *PLAC8 up-regulated* genes (S1 Table).

The most *PLAC8 up-regulated* gene was *ALD5*, coding for a mitochondrial K^+^ activated aldehyde dehydrogenases (ALDH). In yeast, ALDHs have a distinct role in the antioxidant cell responses because they maintain redox balance by supplying reducing equivalents in the form of NADH and NADPH [28]. However, Ald5p seems to play only a minor role as ALDH, because an *ald5* mutant retained 80% of K^+^-activated ALDH activity [29]. ALD4, the major K^+^-activated mitochondrial ALDH, as well as the cytosolic ALD6, were both *PLAC8 down-regulated* genes (S1 Table). Kurita & Nishida [29] showed a more important role of the mitochondrial Ald5p in the regulation or the biosynthesis of electron transport chain components. We measured total cellular respiration in the yeast strain W303-1B transformed with Onzin, FCR1 or the empty vector, and the results (S4 Fig) indicate that the overall oxygen consumption was slightly increased (about 10%) in the PLAC8-expressing strains, irrespective of CdSO_4_ exposure. Overall, the transcriptomic data suggest that the two PLAC8 proteins did not reduce cadmium toxicity and mutagenicity simply by increasing the cell antioxidative response.

### Onzin and FCR1 induce iron-dependent pathways for leucine and iron-sulfur cluster biosynthesis

The *PLAC8 up-regulated* genes included key genes involved in amino acid biosynthesis, such as arginine, histidine, methionine and threonine (S1 Table), but one of the most represented pathways was the super-pathway of leucine, isoleucine, and valine biosynthesis (S2 table, Fig 4). *PLAC8 up-regulated* genes included *ILV2, ILV3, ILV5* and *BAT1*, involved in common reactions of branched-chain amino acids (BCAAs) biosynthesis, and genes specific for the leucine pathway (*LEU1, LEU2, LEU4, LEU9* and *OAC1*). Oac1p, a mitochondrial oxaloacetate transporter, catalyzes the export to the cytoplasm of α-isopropylmalate, an intermediate of leucine biosynthesis produced inside the mitochondrion [30]. Together with *GDH1*, encoding a major enzyme for ammonia assimilation in *S. cerevisiae*, all these *PLAC8 up-regulated* genes are established or potential members of the Leu3p regulon [31], which is transcriptionally regulated by Leu3p. Two *PLAC8 up-regulated* genes of the Leu3p regulon, the acetohydroxy acid reductoisomerase *ILV5* and the BCAA aminotransferase *BAT1* have an additional transcriptional control by Tpk1p, a subunit of yeast protein kinase A, thought to have a role in controlling mitochondrial iron homeostasis [32].

**Fig 4.**
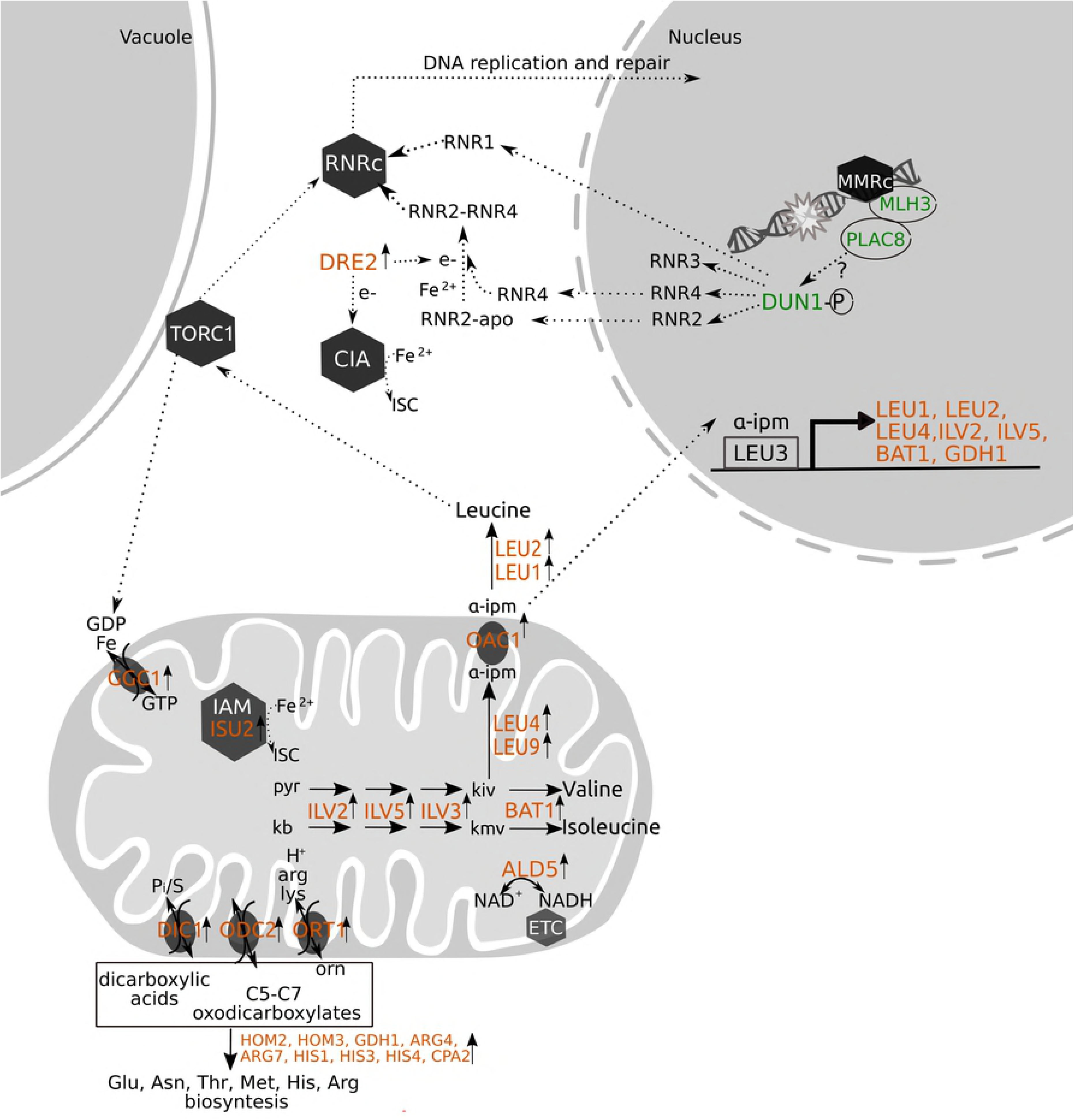
Schematic representation of the cellular functions of the PLAC8 up-regulated genes. PLAC8 up-regulated genes are indicated in orange. Genes for which functional assays were performed are colored in green. Reactions catalyzed by proteins encoded by PLAC8 up-regulated genes are indicated as full arrows. Dotted arrows represent processes or regulatory pathways known from the literature (see text for references). Hexagons represent enzymatic complexes, whereas gray ovals represent membrane carriers. ALD5: Aldehyde Dehydrogenase 5, BAT1: mitochondrial BCAT, CIA: cytosolic ISC assembly, DIC1: Dicarboxylate Carrier 1, DRE2: Fe-S-containing protein supplying reducing equivalents to the early steps of the cytosolic Fe-S assembly (CIA) pathway, DUN1-P: DNA-damage uninducible kinase, in the phosphorilated active form, ETC: electron transfer chain, GDH1: NADP(+)-dependent glutamate dehydrogenase, GGC1: GDP/GTP Carrier 1, IAM: mitochondrial ISC assembly machinery, ILV2: acetolactate synthase, ILV3: dihydroxiacid dehydratase, ILV5: acetohydroxiacid reductoisomerase, ISU2: mitochondrial protein required for iron-sulfur protein synthesis, α-IPM: α-isopropylmalate, pyr: pyruvate, kb: α-ketobutanoate, kiv: α-ketoisovalerate, kmv: a-ketomethylvalerate, LEU1: isopropyl malate isomerase, LEU2: β-IPM dehydrogenase, LEU3: LEUcine biosynthesis transcription factor, acts as an activator in the presence of α-isopropylmalate, LEU4: α-isopropylmalate synthase, LEU9: α-isopropylmalate synthase (paralog of LEU4), MMR: DNA mismatch repair complex, OAC1: OxaloAcetate Carrier 1, ODC2: OxoDicarboxylate Carrier 2, ORT1: ORnithine Transporter 1, RNRc: ribonucleotide reductases complex, TORC1: *t*arget *o*f *r*apamycin complex 1.

The mitochondrion plays a focal role in iron metabolism because is a major generator of heme and iron-sulfur clusters (ISC) cofactors [33]. ISC are among the most ancient and versatile cofactors of proteins involved in many cellular processes such as respiration, DNA synthesis and repair, metabolite biosynthesis, and oxygen transport catalysis [34-37]. Biogenesis of the ISC in Eukaryotes is a highly conserved process that involves the mitochondrial ISC assembly machinery, an export system from the mitochondrion and the cytosolic ISC assembly (CIA) machinery [36]. Notably, *PLAC8 up-regulated* genes included *ISU2* [38] and *DRE2* [39], coding for essential proteins in the mitochondrial and the cytosolic ISC assembly machineries, respectively (S1 Table, Fig 4).

Some PLAC8 up-regulated components of the BCAA biosynthetic pathway can influence ISC cofactors biosynthesis in yeast. For example, mitochondrial ISC biosynthesis is regulated through Leu1p, an abundant cytoplasmic ISC-containing enzyme [40] and a key regulator in the mitochondrial-cytoplasmic ISC balance [41]. Leu1p shares high homology to the iron regulatory protein Irp1 in mammalian cells, suggesting an influence on iron metabolism within the cell [42]. Iron deficiency is thought to influence the pathway of leucine biosynthesis by reducing the activities of multiple ISC-containing enzymes, including Leu1p and Ilv3p [43].

Overall, the transcriptomic data clearly indicate, in cadmium-exposed yeast expressing both PLAC8 proteins, the up-regulation of leucine and ISC biosynthesis, two iron-dependent pathways that involve the mitochondrion.

### Expression of both PLAC8 proteins in yeast does not modify intracellular iron content

A correlation between iron homeostasis and cadmium response has been revealed by genome-wide screening of *S. cerevisiae* deletion mutant collections, as many cadmium-sensitive mutants were affected in genes related to iron homeostasis [44, 45]. Cadmium interferes with iron homeostasis by reducing iron uptake, since iron addition rescued cadmium-sensitivity of yeast mutants [44] and increased cadmium tolerance of *S. cerevisiae* [46]. Moreover, cadmium exposure stimulated the expression of several yeast genes related to iron uptake [47].

The “iron regulon” is a group of ∼30 genes mostly involved in iron acquisition, activated upon iron deficiency by the iron-sensing transcription factors Aft1p and Aft2p [43]. Several *PLAC8 down-regulated* genes (S1 Table) are known members of the yeast iron regulon, like the high affinity iron uptake system (*FET3* and *FTR1*), components of the siderophore transport system (*FIT2* and *FIT3, SIT1*, alias *ARN3*), and the mRNA-binding protein *TIS11* (alias *CTH2*). Their expression pattern indicates an activation of the iron regulon in cadmium-exposed yeast cells expressing the empty vector, thus suggesting that they experience a condition of iron depletion, as compared with PLAC8-expressing yeasts. We therefore measured total iron content in yeast exposed to cadmium and expressing FCR1, Onzin or the empty vector. No statistical differences were found in the total iron content of yeast cells exposed for 8 h to 25 μM CdSO_4_, the same conditions used for the transcriptomic experiment, independently on the intracellular Cd-concentration (S3 Table). Although it is still possible that different cell compartments may experience different iron concentrations, these results indicate that the expression pattern of the iron regulon genes in the PLAC8-expressing yeast does not simply reflect an increased iron content. Chen and colleagues [48] demonstrated that inhibition of ISC biosynthesis induced the iron regulon in spite of high cytosolic iron levels. Ueta and colleagues [49] later showed that an ISC-dependent interaction between monothiol glutaredoxins and Aft1p in *S. cerevisiae* specifically promotes the dissociation of Aft1p from its target promoters under iron sufficiency. Thus, we may speculate that the up-regulated expression of ISC biosynthesis in the two PLAC8-expressing yeast strains may similarly downregulate transcription of the iron regulon.

### Phenotype of PLAC8-expressing yeast: suggestions from the transcriptome

Although the phenotype of Onzin and FCR1-expressing yeast exposed to cadmium included increased growth (Fig 2) and reduced DNA mutation rate (Table 1), no genes specifically related to cell cycle control or DNA damage repair could be identified among the *PLAC8-regulated* transcripts. Of course, we cannot exclude a post-transcriptional regulation of these pathways, but we could also speculate on other possible scenarios based on the transcriptomic data.

The main pathway up-regulated by both PLAC8 proteins in cadmium-exposed yeast was the leucine biosynthetic pathway. This pathway generates leucine, one of the most conserved and potent *TORC1* (*t*arget *o*f *r*apamycin complex 1) activating growth signals (Fig 4). TORC1 is a protein complex, conserved throughout Eukaryotes, that functions as a master regulator of cell proliferation, survival and growth [50]. The mitochondrial branched amino acid transferase Bat1p and the β-isopropylmalate dehydrogenase Leu2p, both *PLAC8 up-regulated* genes, can reversibly metabolize respectively leucine or β-isopropylmalate to α-ketoisocaproate (KIC), another effective *TORC1* activator [51]. A mitochondrial GTP/GDP transporter (GGC1) up-regulated by both PLAC8 proteins has been also characterized as a component of the rapamycin/target of rapamycin (TOR) signaling pathway [52]. Lesuisse and colleagues [53] demonstrated that GGC1 mutants accumulated iron in the mithocondria and suggested a role in intracellular iron balance, either by directly transporting iron from mitochondria to cytoplasm as heme, Fe–S or other forms, or by transporting molecules that influence iron uptake and distribution. In yeast, TORC1 regulates cellular responses to a variety of environmental stresses and is a determinant of cell survival in response to DNA damage thanks to increased dNTPs synthesis by ribonucleotide reductases (RNR, [54]).

Several proteins involved in DNA synthesis and repair contain ISC cofactors, including replicative DNA polymerases and primase, DNA helicases, nucleases, glycoylases and demethylases [55, 56]. An interesting mechanism proposed by Arnold and colleagues [57], named *DNA charge transport*, suggests that DNA processing enzymes containing the ISC cofactor may use electrons released from redox active iron to rapidly and efficiently scan DNA over long molecular distances for mismatches and damages. It is therefore not surprising that impairments in the mitochondrial or the cytosolic ISC assembly machineries are connected with nuclear genomic instability [58, 59]. Therefore, we could speculate that, by up-regulating ISC biosynthesis, yeast cells expressing the two PLAC8 proteins may increase their survival because they provide essential cofactors to enzymes involved in DNA damage repair, thus reducing the high DNA mutation rate observed in cadmium-exposed yeast cells (Table 1). MM19, a late-acting CIA component likely implicated in the delivery of ISC into nuclear and cytosolic apoproteins [56], was found to be necessary to *Schizosaccharomyces pombe* to grow on cadmium [60].

Although these hypotheses are only based on transcriptomic data, some experimental evidence suggests a link between PLAC8 protein functions in cadmium tolerance, ISC biosynthesis and DNA damage repair. The nuclear DNA damage caused by defects of ISC biosynthesis activates at least two different signalling pathways that converge at Dun1p, a protein kinase that controls the DNA damage response in yeast [61]. The DNA damage checkpoint mediated by the Mec1p–Chk1p– Dun1p signaling transduction pathway was found to be activated by dysfunctions in ISC-targeting factors, which are not required for the biogenesis of ISC but act specifically for transferring ISC to mitochondrial target apoproteins [62]. By contrast, dysfunctions of the core mitochondrial ISC assembly machinery induced a second pathway involving a Mec1p-independent activation of Dun1p. Similarly, Sanvisens and colleagues [63] found that depletion of core components of the mitochondrial ISC assembly activated a Dun1p-dependent but Mec1p- and Rad53p-independent pathway leading to increased dNTPs biosynthesis, needed for DNA repair.

Thus, Dun1p is a central actor in the activation of the DNA damage checkpoint induced by dysfunctions of the ISC assembly machineries [62]. Abbà and colleagues [19] found that Dun1p was necessary for the cadmium tolerant phenotype of FCR1 in a Mec1p-independent pathway. Here, we show that Dun1p was also required for the cadmium tolerant phenotype of Onzin-expressing yeast (Fig 5), suggesting that both PLAC8 proteins may activate a specific Dun1p-dependent DNA damage checkpoint pathway similar to the one described by Pijuan et al, [62] and Sanvisens et al, [63]. Indeed, Dun1p activates the RNR at multiple levels (i.e. phosphorylation of the Sml1p RNR1-ihnibitor; phosphorylation and degradation of Dif1p, promoting the redistribution of the small RNR2 and RNR4 subunits to the cytoplasm where the catalytic RNR1 subunit resides; phosphorylation of the Crt1 transcriptional RNR2/3/4 repressor; Fig 5, [56]). Interestingly, the *PLAC8 down-regulated* mRNA-binding protein TIS11, belonging to the iron regulon, targets for degradation specific mRNAs encoding proteins that contain iron as a cofactor, including the small subunit of the ribonucleotide reductases (RNR2) as part of metabolic remodeling to conserve and optimize iron utilization [64].

**Fig 5.**
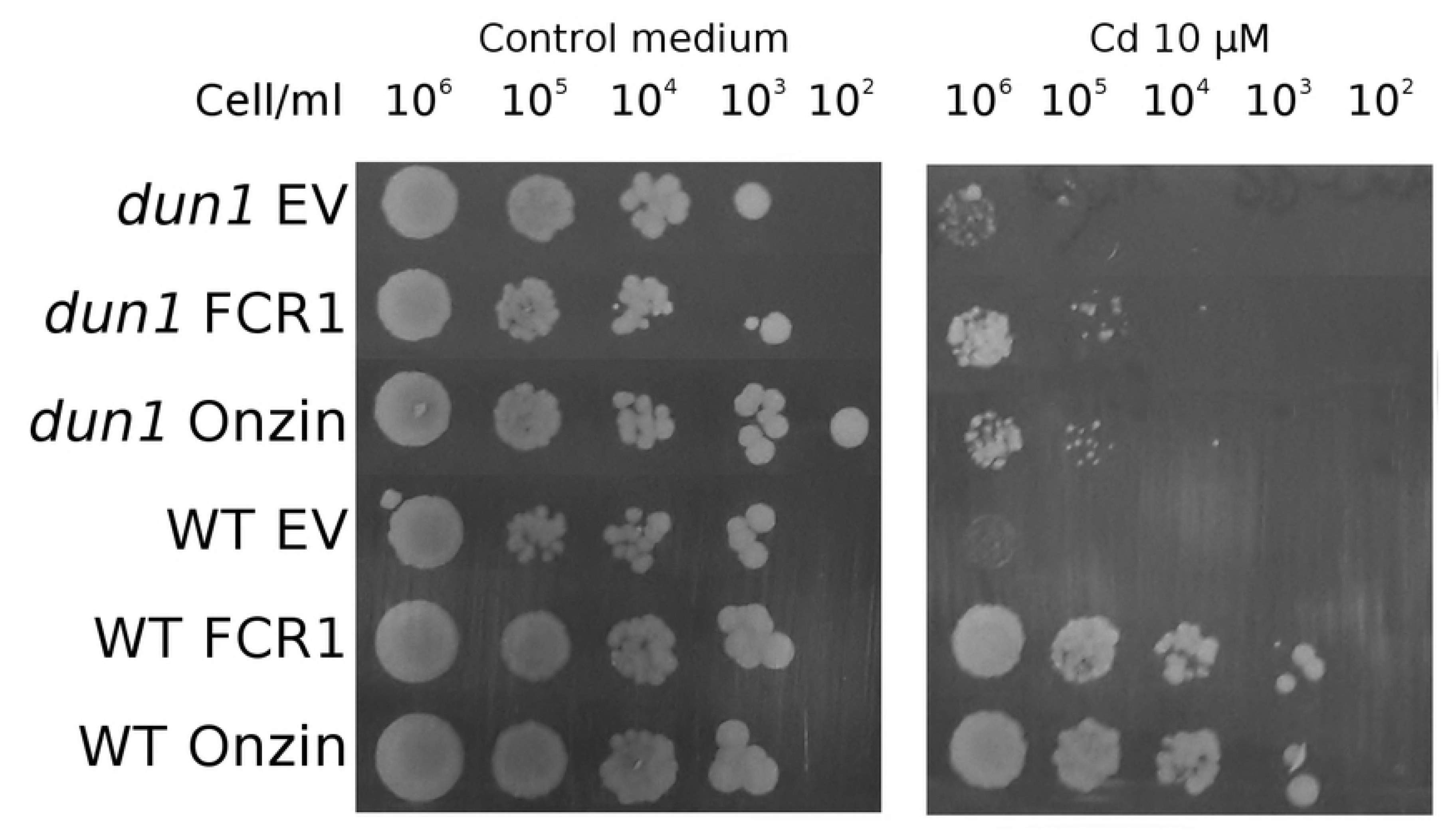
Influence of DUN1 deletion on cadmium tolerance in yeast cells expressing FCR1 and Onzin. Spot dilution assay of wild-type W303 (WT) and *dun1* mutant strains expressing FCR1, Onzin or the empty vector pFL61 (EV). Strains were plated in ten-fold serial dilutions onto YNB-D medium, with or without 10 µM CdSO_4_.

### Ancestral functions of PLAC8-containing proteins?

Proteins containing the PLAC8 domain are widely distributed in Eukaryotes and the PLAC8 proteins characterized so far are assigned two alternative functions, either in heavy metal resistance or as cell growth regulators. In this work, we showed that Onzin, the reference for PLAC8 cell growth regulators [2], could induce cadmium resistance in *S. cerevisiae*, demonstrating that these two functions are not mutually exclusive in PLAC8 proteins. Another example of how the same PLAC8 protein can display both functions is the rice OsPCR1 that, even though with different acronyms, has been reported as cell growth regulator [16] and able to confer cadmium-resistance [65].

Although the *S. cerevisiae* genome does not encode PLAC8 proteins, expression of FCR1 and Onzin induced ancient and highly conserved pathways that play a central role in cell growth and metabolism. In addition to iron homeostasis, components of the leucine biosynthetic pathway are involved in additional regulatory functions related to general cell growth and metabolism. The mitochondrial BCAA transaminase Bat1p is particularly interesting because it shows striking sequence similarity to the mammalian protein Eca39, a known target for c-Myc regulation [66, 67]. Myc proteins are involved in cell proliferation and differentiation in vertebrates [68] and although they have not been found in yeast, a shortened G1 stage was observed in *bat1* yeast mutants [69]. Thus, in addition to a role in BCAA biosynthesis and ISC translocation, Bat1p is also involved in cell cycle regulation. It is intriguing that Onzin, another known target of c-Myc activity in mammals [2, 70], increased cell survival and growth in the presence of cadmium by modulating the same conserved biosynthetic pathways.

Leucine biosynthesis was the main pathway up-regulated by both PLAC8 proteins in cadmium-exposed yeast. In addition to its interaction with iron homeostasis, this pathway generates the amino acid leucine, one of the most conserved TORC1 activating growth signals. TORC1 is a master regulator of cell proliferation, survival and growth found in yeast and mammals [50] and is activated through a pathway, partly conserved in Eukaryotes, that uses leucyl-tRNA synthetases as leucine sensors [71]. In yeast, this pathway regulates cellular responses to a variety of environmental stresses and is a determinant of cell survival in response to DNA damage, thanks to increased dNTPs synthesis [54]. It would be interesting to understand whether the PLAC8 and the TORC1 induced pathways may potentially interact in yeast.

In conclusion, our data clearly demonstrate that two PLAC8 proteins with different described functions in taxonomically distant organisms induced the same phenotype when expressed in *S. cerevisiae*, suggesting a common ancestral function. Having identified some processes and pathways regulated by both PLAC8 proteins, the two main functions ascribed to these proteins (i.e. increased cadmium tolerance and regulation of cell proliferation) appear more related, as they could be both linked to leucine and ISC biosynthesis. Although this hypothesis requires further investigations, our results open new perspectives on the role of the PLAC8 protein domain in Eukaryotes and provide guidelines to explore in more details the exact role of the PLAC8 domain in the activation of important biological processes that ensure nuclear DNA integrity during cell division and metal stress responses.

## Methods

### Yeast strains and growth conditions

All yeast strains used in this work are listed in supplementary the S1 Appendix. EAY1269 wild type strain was kindly provided by Prof. Eric Alani (Cornell University, Ithaca, NY, USA). Yeast strains were grown at 30°C on Yeast Extract Peptone medium (YP) supplemented with 2% (W/v) glucose (YPD). Yeasts transformed with episomal plasmids were grown at 30°C on Yeast Nitrogen Base (YNB) medium supplemented with essential amino acids and either 2% (W/v) glucose (YNB-D) or 2% (W/v) galactose (YNB-Gal). All reagents were purchased from Sigma-Aldrich. Yeasts were transformed according to Gietz & Woods [72]. Transformation was confirmed by colony-PCR as described by Sambrook & Russell [73] with FL1 and FL2 primers (S2 Appendix). Strains transformed with pFL61-derived vectors were grown in medium lacking uracil. Yeasts co-transformed with pMS207 vector were grown in medium lacking leucine also. Media and growth conditions for Yeast Two Hybrid were prepared as previously described [19].

### Protein sequences alignment

The protein alignment was performed using the Phylogeny.fr platform [74]. Sequences were aligned with MUSCLE (v3.8.31) configured for highest accuracy (MUSCLE with default settings).

### Spot dilution assay

Spot dilution assays were used to monitor cell growth at various cadmium concentrations and were conducted as previously described [19], with the following modifications: for all conditions, overnight yeast cultures were diluted to OD_600_=0.1 and cultured at 30°C until OD_600_=0.2. Subsequently, 4 μl of serial dilutions (ranging from 5×10^7^ to 5×10^2^ cells/ml) of each strain were spotted onto control or cadmium-amended YNB-D medium and incubated at 30°C for 4 days.

### Determination of cadmium half inhibitory concentration (IC50)

One clone of each EAY1269 yeast strain (transformed with the empty vector or expressing FCR1 or Onzin or Onzin^Δ28-38^) was inoculated in 5 ml YNB-D medium and incubated at 30°C overnight. The following morning, each pre-inoculum was diluted in fresh YNB-D medium at optical density OD_600_=0.1, and poured in 96-well plates. The cultures were amended with increasing concentrations CdSO_4_ (from 0 to 400 µM), incubated at 30° C and 150 rpm, and the OD_600_ was measured by using a microplate reader (TECAN) after 24 hours. The experiment was independently replicated 3 times, with 5 technical replicates per concentration and per clone (n=15). IC50 values, representing the Cd concentration causing a 50% of inhibition of the yeast growth, were calculated using the ED50 Plus v1.0 available on line, and previously utilized and validated [75]. S1 Fig shows the distribution of the data from the three experiments, with standard deviation. Different letters indicated statistically different results (p<0.05). Single data from the three experiments, Shapiro Wilk test for normal distribution, p-values calculated by applying the ANOVA with Tukey as post-hoc test are are available in the S1 Data.

### Growth curve and cell viability assay

For the growth curve assay, three clones of each EAY1269 yeast strain (transformed with the empty vector or expressing FCR1 or Onzin) were inoculated in 5 ml YNB-D medium and incubated at 30°C overnight. The following morning, each pre-inoculum was diluted to 50 mL in fresh YNB-D medium at optical density OD_600_ =0.1, and split into two sterile flasks, one amended with 25 µM cadmium sulphate and the other left unamended. The optical density of cadmium-treated and control cultures was measured after 2, 4, 6, 8, 10, and 24h of incubation at 30°C. The data were tested for normal distribution with the Shapiro Wilk test (p-value > 0.05) and the ANOVA was applied with Tukey as post-hoc test. Single data and the calculated p-value for data at 6 and 8 hours are shown in the S2 Data. Fig 2B shows the result of a single experiment (n=3) that is representative of three independently repeated experiments, each one based on n=3 or n=5 biological replicates, and the raw data obtained after 8hrs of incubation are reported in the S3 Appendix. All the statistics were performed using PAST [76].

To evaluate the number of viable yeast cells during growth in liquid culture, the number of Colony Forming Units (CFU) was measured by plating 100 μL of the appropriate dilutions (based on the initial OD) onto standard YNB-D medium after 6, 8, 10 and 24 h of growth in control or cadmium-amended liquid YNB-D. Fig 2C reports the distribution of six biological replicates for the time-points 6, 10 and 24 h and three biological replicates for the 8h time-point, obtained in two independent experimental repetitions. Single data, Shapiro Wilk test for normal distribution, p-values calculated by applying the ANOVA with Tukey as post-hoc test are reported in the S3 Data.

### Onzin coding sequence isolation and expression in yeast

All oligonucleotide sequences are listed in S2 Appendix. cDNA corresponding to *Mus musculus* Onzin CDS was synthesized with Qiagen OneStep RT-PCR Kit (Qiagen, Venlo, Netherlands) using total RNA from mouse blood and Not_Onzin1f/Not_Onzin2r primers which carry *NotI* restriction site at the 5’ end. PCR program was as follows: 60s at 98°C for 1 cycle; 10s at 94°C, 30s at 60°C, 40s at 72°C for 35 cycles; 10 min at 72° C for 1 cycle. Amplified DNA was digested with *NotI* restriction enzyme and cloned into the pFL61 Vector. Sequence as well as direction of the inserted gene were assessed by PCR and sequencing using primers FL1 and Not_Onzin2r. This vector was named pFL61-Onzin. The mutant allele Onzin^Δ28-38^ was obtained using a two-round mutagenesis PCR. In the first round, the 5’ end of the sequence was amplified with primers Not_Onzin1f/Onzindel_r while the 3’ end was amplified using primers Onzindel_f/Not_Onzin2r using Thermo Phusion High Fidelity DNA Polymerase (Thermo, Waltham, MA, USA). PCR program was as follows: 30s at 98°C for 1 cycle; 10s at 98°C, 30s at 62°C, 20s at 72°C for 35 cycles; 10 min at 72° C for 1 cycle. In the second round of PCR, the two purified products from the first amplification where used as template for a fusion-PCR using primers Not_Onzin1f /Not_Onzin2r and the same enzyme used before. PCR conditions were as follows: 30s at 98°C for 1 cycle; 10s at 98°C, 40s at 62°C, 20s at 72°C for 35 cycles; 10 min at 72° C. The PCR product was then digested with the restriction enzyme *NotI*, ligated into the plasmid pFL61 previously digested with the same enzyme and cloned into *E. coli*. Transformed colonies were screened by colony PCR with primers FL1/Not_Onzin2r and positive plasmids were sequenced with primers FL1 and FL2 to confirm the construct sequence. The pFL61-FCR1 construct has been obtained in a previous work [19].

### Synthesis of EGFP-tagged Onzin construct and microscopy observations

Construction of C-terminal Enhanced Green Fluorescent Protein (EGFP) tag of Onzin was performed by amplifying the Onzin cDNA with primers HindIII_Onzin_f/XmaI_Onzin_r, carrying *HindIII* and *XmaI* restriction sites and by removing the CDS stop codon. The DNA obtained was then digested and directionally ligated in a previously digested (with the same enzymes) pEGFP-N1 vector and cloned into *E. coli*. Colony PCR with primers HindIII_Onzin_f/Not_EGFP_r was used to identify and purify the right construct which was then confirmed by sequencing with the same primers. To express Onzin-EGFP gene in yeast cells, the purified construct was used as template for a second PCR with primers Not_Onzin1f/Not_EGFP_r and the PCR product was digested with *NotI* enzyme, ligated in pFL61 plasmid previously digested with the same enzyme and cloned in *E. coli*. Colony PCR using primers Not_Onzin1f / FL2 was used to confirm size and 5’-3’ orientation of the construct. This vector was called pFL61-Onzin-EGFP. The pFL61-FCR1-EGFP construct has been obtained in a previous work [19].

Yeast cells expressing an inducible tomato protein carrying a nuclear localization signal (NLS)-RFP (plasmid pMS207, courtesy of prof. Maya Schouldiner, Weizmann Institute, Rehovot, Israel) were transformed either with the plasmid carrying the constitutive Onzin-EGFP tagged construct (pFL61-Onzin-EGFP) or with the plasmid carrying the FCR1-EGFP tagged construct. Double transformants were grown overnight with galactose as the sole carbon source (YNB-Gal). The localization of EGFP-tagged and RFP-tagged proteins was observed on a Leica TCS SP2 confocal microscope, using a long-distance 40X water-immersion objective (HCX Apo 0.80). For EGFP visualization, an Argon laser band of 488nm was used for excitation and the emission window was recorded between 500 and 525nm, while for the RFP-tagged protein a laser light of 554 nm was used, and an emission window of 581 nm.

### Yeast-Two-Hybrid

The yeast two-hybrid assay was performed using the DupLEX-A yeast system (Origene Technologies, Rockville, MD, USA) as described in previous work [19]. The MmOnzin coding sequence was cloned in frame with the DNA binding domain of LexA into the pEG202 vector using primers Eco_MmOnzin_f/Not_Onzin2r. The C-terminal region of *MLH3* from *Mus musculus* was isolated with Qiagen OneStep RT-PCR Kit (Qiagen, Venlo, The Netherlands) using total RNA from mouse blood and primers Eco_MmMLH3_f/Eco_MmMLH3_r (S2 Appendix) then cloned downstream in frame with the activator domain of B42 into the pEG202 vector. Sequence and orientation were confirmed by colony-PCR and Sanger sequencing. All the other vectors and strains used in the yeast-two-hybrid assay were generated in previous work [19].

### Lys+ reversion assay

EAY1269 strains (transformed with the empty vector or expressing FCR1 or Onzin) were analyzed for reversion to the Lys^+^ phenotype according to Tran et al, [25]. Cells were cultured in YNB-D for 24 hrs at 30°C and 150 rpm shaking with or without 1μM CdSO_4_, a concentration that did not influence cell growth in 24 hrs. At the end of the incubation, 100 μl of the appropriate dilution were plated on normal and Lys^-^ dropout medium. Colonies were counted after four days. At least n=20 independent biological replicates for each strain and condition were analyzed (reported in Table 1). Reversion rates were determined as previously described [77]. Confidence intervals of 95% were determined as described by [78]. Each median reversion rate was normalized to the empty vector median rate to calculate the fold increase in mutation rate.

### RNAseq analysis

EAY1269 cells expressing Onzin, FCR1 or transformed with the empty vector (pFL61) were grown in three biological replicates (n=3), as described for the growth curve, on YNB-D with 25 μM CdSO_4_. After 8hrs, cells were harvested by centrifugation, washed with sterile water and frozen in liquid nitrogen. Total RNA was extracted in a CTAB-based extraction buffer (2% CTAB, 2% PVP, 100 mM Tris-HCl pH 8, 25 mM EDTA, 2 M NaCl, 2% β-mercaptoethanol, 1% (W/v) PVPP). The homogenates were incubated 5 min at 65°C, extracted twice in chloroform:isoamyl alcohol 24:1 (v/v), precipitated with an equal volume of LiCl 10M (on-ice over-night precipitation), resuspended in an SSTE buffer (1 M NaCl, 0.5% (W/v) SDS, 10 mM Tris-HCl pH 8, 1 mM EDTA), extracted with phenol:chloroform:isoamyl alcohol 25:24:1 (v/v/v), extracted in chloroform:isoamyl alcohol 24:1 (v/v), precipitated with 100% ethanol (2 hrs, -20°C), washed in 80% ethanol and resuspended in DEPC-treated water. The quantity and quality of the extracted RNA was evaluated with a 2100 Bioanalyzer (Agilent Technologies, Santa Clara, CA, USA). The mRNA was sequenced by Illumina technology with Hiseq 2000, raw reads of 50 NTs were firstly assessed for their quality using FastQC and then mapped on *S. cerevisiae* S288C genome with Burrows-Wheeler Aligner [79]. Resulting BAM files were processed using SAMtools [80]. Only reads with phred score greater than 15 were considered in the analysis. The genome sequence and annotation files were obtained from SGD database (http://www.yeastgenome.org). The differential gene expression was assessed with DESeq package [81], retaining only genes with FDR-corrected p-value <0.1. Genes with read counts equal to zero in all replicates of at least one experimental condition were excluded from the analysis. The fold change was calculated with respect to the empty-vector expressing strain and genes with log_2_ fold-change >1 or <-1 and with adjusted p-value <0.05 were considered respectively up-or down-regulated by the PLAC8 proteins. Gene Ontology and pathway enrichment were evaluated using DAVID Bioinformatics Resources 6.8 and Saccharomyces Genome Database [82, 83].

### Respiratory activity

Yeast cells were cultured over night at 28°C in YNB medium supplemented with 0.6% glucose, then cells were treated for 4 hrs with up to 200µM CdSO_4_. Oxygen consumption rate was measured at 30°C using a Clark-type oxygen electrode (Oxygraph System Hansatech Instruments England) with 1 mL of air-saturated respiration buffer (0.1 M phthalate–KOH, pH 5.0), 0.5% glucose. The experiment was independently performed twice (n=2), with one biological replicate per sample.

### Metal content analysis

The EAY1269 yeast strains, transformed with the empty vector or expressing either FCR1 or Onzin, were inoculated in 5 ml YNB-D medium and incubated at 30°C overnight. The following morning, each pre-inoculum was diluted to 50 mL of the control YNB-D medium, or in the same medium supplemented with 25 μM CdSO_4_. Five different clones were used for each transformant, per each experimental condition (n=5). Yeast cultures were harvested after 8h and centrifuged at 4000 rpm for 5 min. Cells were washed three times with 10 mM EDTA in 50 mM Tris–HCl buffer (pH 6.5), and with milli-Q water. Finally, samples were dried at 60 °C for 2 days, and subsequently mineralized with 1 ml HNO_3_ 6M in a bath at 90°C for 1h. After dilution to a final concentration of 1M HNO_3_, the metal content was determined using Induced Coupled Plasma (ICP-OES Optima 7000 DV, Perkin Elmer). Controls made up of milli-Q water and nitric acid 1M. Statistics have been performed by using the Past software [76]. Normality of data was assessed by Shapiro Wilk test. Pairwise differences have been calculated by the Mann-Whitney test for differences in the medians. Different letters indicate significant differences between samples (p<0.05). The raw data and single p-values are reported in the S4 Data.

### Data Availability

All relevant data are within the manuscript and its Supporting Information files. The RNAseq data from this publication have been deposited to the SRA database, have been assigned the identifier SRP145576 and will be available after acceptance.

## Acknowledgements

We thank Simona Abbà (Institute for Sustainable Plant Protection of the National Research Council of Italy, IPSP-CNR, Turin, Italy) for Onzin gene isolation and support through the project, Eric Alani (Cornell University, Ithaca, NY, USA) for providing the EAY1269 strain and technical support with the Lys2+ reversion assay, and Maya Schuldiner (Weizmann Institute of Sciences, Rehovot, Israel) for the pMS207 vector and technical support in the FCR1 and Onzin cellular localization.

## Supporting information captions

**S1 Fig. Half Inhibitory Concentration (IC50) of CdSO**_4_.

Yeast expressing FCR1, wild-type Onzin, truncated Onzin^Δ28-38^ or the empty vector pFL61 (EV) were exposed to increasing Cd concentrations and IC50 have been determined. The distribution of the data from three independent experiments is shown in the figure. The square symbol indicates the mean value. The wiskers indicate the minimum and the maximum values. The top and the bottom of the rectangle indicate ± standard deviation, while the central line of the rectangle indicates the 50%. Statistically different results are indicated with different letters (P<0.05 by ANOVA with Tukey as post-hoc test, Shapiro Wilk as normality test).

**S2 Fig. Yeast-Two-Hybrid to investigate FCR1 and Onzin interactions with different Mlh3 proteins.** Genes coding for Mlh3p were isolated from *S. cerevisiae* (ScMlh3) from *O. maius* (OmMlh3) and from *Mus musculus* (MmMlh3). Yeasts were plated with ten-fold dilutions onto galactose and β-galactosidase containing-medium lacking leucine. The dark blue color of FCR1/ScMlh3 and FCR1/OmMlh3 colonies indicates a strong protein–protein interaction. Onzin is able to interact with both ScMlh3 and MmMlh3, but the blue color is less intense. No self-activation was observed when cells were co-transformed with a vector expressing an irrelevant protein such as *O. maius* superoxide dismutase (OmSOD1), that was used as negative control. A and B represent two indipendent replicates of the same experiment.

**S3 Fig. Genes differentially regulated by PLAC8 proteins in the transcriptome of yeast cells exposed to cadmium.** The diagram shows the number of genes differentially regulated by Onzin (Onzin-regulated), by FCR1 (FCR1-regulated) or by both proteins (PLAC8 regulated) in yeast cultures grown for 8 h in cadmium-containing medium (25 µM), as compared to the yeast strain transformed with the empty vector.

**S4 Fig. Oxygen consumption rate of yeast strains expressing FCR1, Onzin or the empty vector.** Yeast cells were exposed to different CdSO_4_ concentrations, ranging from 0 to 200 µM. The data from two independent assays are plotted.

**S1 Table. Regulation of gene expression in yeast cells exposed to cadmium and expressing FCR1 or Onzin.** Sheet A: FCR1 up-regulated yeast genes; sheet B: Onzin up-regulated yeast genes; sheet C: FCR1 down-regulated yeast genes; sheet D: Onzin down-regulated yeast genes. PLAC8-regulated genes are highlighted in red. The fold change was calculated with respect to the empty-vector expressing strain and genes with log_2_ fold-change >1 or <-1 and with adjusted p-value <0.05 were considered respectively up- or down-regulated.

**S2 Table. GO categories and KEGG pathways enriched in PLAC8-regulated genes.** Categories enriched with p-value<0.05 are reported.

**S3 Table. Iron and cadmium content in Cd-treated yeasts expressing Onzin, FCR1 or the empty vector.** Intracellular content of iron and cadmium in yeast cells exposed to cadmium (25 µM for 8h) was measured after total biomass digestion by ICP-OES. Different letters indicate significant differences between samples (p<0.05 according to Mann-Whitney test for differences in the medians with n=5).

**S1 Appendix. Yeast strains used in this work.**

**S2 Appendix. Yeast growth of unexposed and cadmium-exposed (Cd) cell cultures.** The optical density (OD 600nm) of yeast cultures expressing FCR1, Onzin and the empty vector pFL61 (EV) was measured after growth in control medium or in Cd-containing medium (25uM) for 8 hrs at 30°C, 150 rpm. The table reports the raw data for three independent experiments, each one including n=3 or n=5 biological replicate of each sample.

**S3 Appendix. List of oligonucleotides used in this work.**

**S1 Data. Source data and detailed statistics for S1 Fig.**

**S2 Data. Source data and detailed statistics for Fig 2B.**

**S3 Data. Source data and detailed statistics for Fig 2C.**

**S4 Data. Source data and detailed statistics for S3 Table.**

